# Height-diameter allometry for tropical forest in northern Amazonia

**DOI:** 10.1101/2021.08.04.455098

**Authors:** Robson Borges de Lima, Eric Bastos Görgens, Fernando Elias, Jadson Coelho de Abreu, Aldine Luiza Baia, Cinthia Pereira de Oliveira, Diego Armando Silva da Silva, Anderson Pedro Bernardina Batista, Robson Carmo Lima, Eleneide Doff Sotta

## Abstract

Height measurements are essential to manage and monitor forest biomass and carbon stocks. However, accurate estimation of this variable in tropical ecosystems is still difficult due to species heterogeneity and environmental variability. In this article, we compare and discuss six nonlinear allometric models parameterized at different scales (local, regional and pantropical). We also evaluate the height measurements obtained in the field by the hypsometer when compared with the true tree height. We used a dataset composed of 180 harvested trees in two distinct areas located in the Amapá State. The functional form of the Weibull model was the best local model, showing similar performance to the pantropical model. The inaccuracy detected in the hypsometer estimates reinforces the importance of incorporating new technologies in measuring individual tree heights. Establishing accurate allometric models requires knowledge of ecophysiological and environmental processes that govern vegetation dynamics and tree height growth. It is essential to investigate the influence of different species and ecological gradients on the diameter/height ratio.

## Introduction

The height of the tree at the species and individual level is very relevant information for understanding the autecological characteristics (e.g., fitness, size, ontogenetic stage, trophic position) and essential for monitoring biomass and carbon stocks [1]. Tropical humid forests are formed by a high diversity of species, a complex vertical structure, a dense canopy and high relative humidity. These characteristics make height measurements difficult, inefficient and with a high error [2,3]. As a result, biometrists and forest ecologists tend to measure only the diameter of the trees and estimate the height using allometric models.

Classic logarithmic or polynomial models are often used for non-linear height-diameter relationships [4,5]. Some studies have evaluated the potential of alternative fitted models on a larger scale (continental and pan-tropical) based on the inclusion of environmental variables intrinsic to each region of the tropics [6,7]. Studies using the power-law model [6,8,9] and the Weibull model demonstrate a considerable reduction in errors in height estimates when fitted for local-scale [1,3,10,11]. In addition, approaches with monomolecular and hyperbolic models based on diameter growth [12], and including spatial information [13,14], resulted in high precision using a reduced set of predictor variables.

Several efforts have been made to develop height-diameter allometric models for tropical forests at continental scales in Africa [3] and Asia [15]. In South America, we noticed several proposals for models at regional scales [6,7,16,17]. However, there are still important gaps related to height-diameter allometric modelling for the different Amazonian humid tropical forest regions. The available field data covers less than 0.0013% [18] and are concentrated along the great rivers and highways. As a result, 42% of the Brazilian Amazon is more than 50 km far away from a forest inventory plot [18– 20].

One of the most unknown regions of the Brazilian Amazon is the northeast region of the Amazon (northern Pará and the Amapá States). Such regions demand studies to understand the main implications of using models from the literature. It is essential to provide a better understanding of the variation in biomass and carbon stocks [8,21], as well as producing responses regarding the transferability of models from one region to another. The objective of this paper is to develop local height-diameter allometric models for the tropical humid forest in the northeast of the Brazilian Amazon. We also look to compare the developed local model to the main pan-tropical and regional models from the literature [6,16,17]. Finally, we try to understand the accuracy in estimating the tree height using allometric models, hypsometer and reference height.

## Materials and Methods

### Study area

The data were recorded from two sites in the State Forest of Amapá -FLOTA / AP (Fig 1). The first site is located between the municipalities of Calçoene and Oiapoque, (2º57’16.00’’ N and 51º27’57.59’’ W), accessed via the BR-156 and the Caciporé River, 630 km from Macapá. The second site is located in the municipality of Porto Grande (00°34’55,7’’ N and 52° 03’54,9’’ W), 130 km from Macapá.

**Fig 1.**
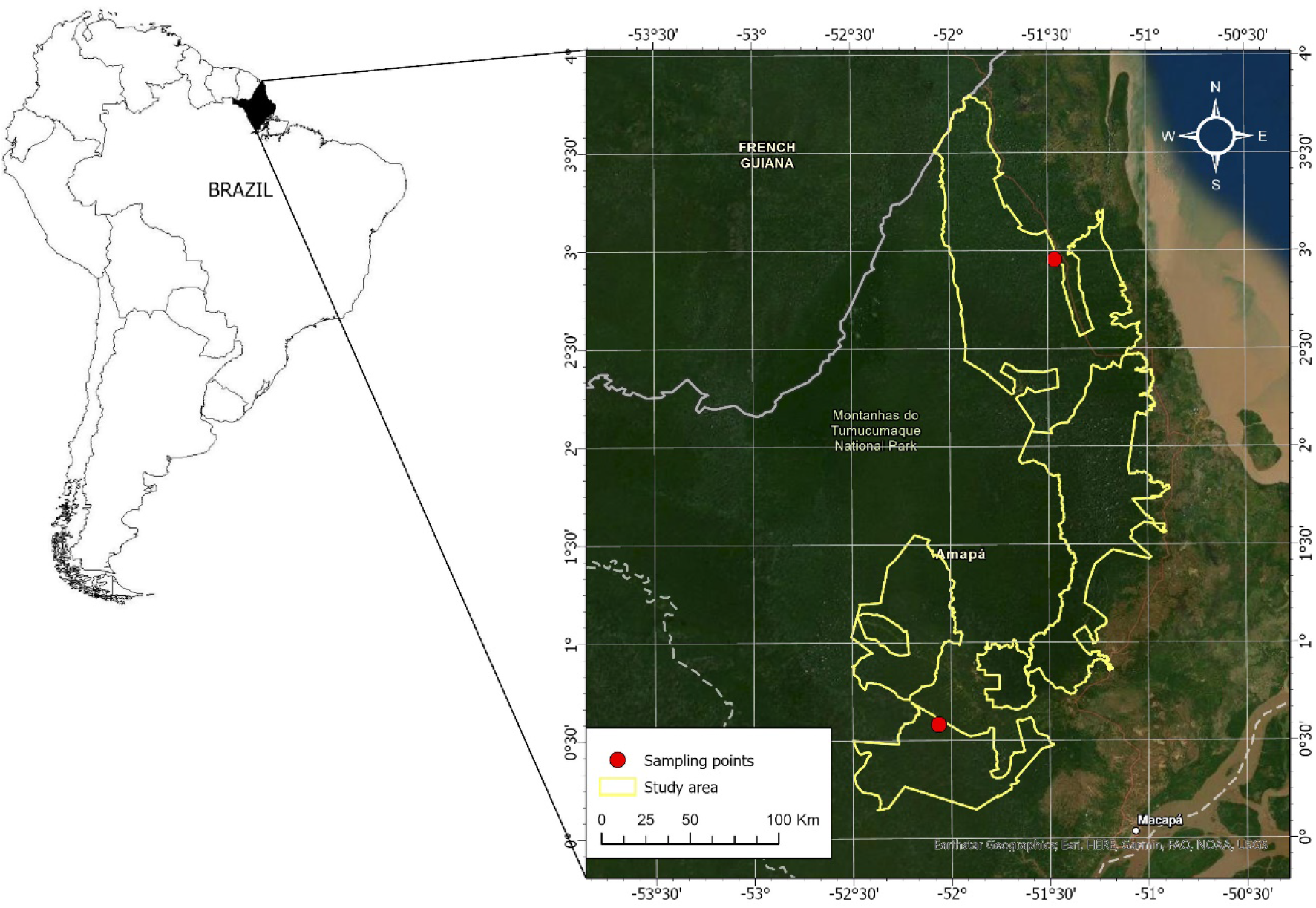
Study locations: area A located in Calçoene and Oiapoque municipalities (A) and area B located in Porto Grande municipality (B).

The climate is classified according to Koppen as Amw (monsoon tropical climate with summer-autumn rains). The rainy season goes from December to June and with a relatively dry period from July to November [22]. The average annual rainfall (last 10 years) is approximately 3,225 ± 138 mm, with September being the shortest rainy and May showing the highest rainfall. Both sites have low thermal amplitude, varying from 24.8 ± 0.15 ° C during the coldest month to 26.9 ± 0.10 ° C during the hottest month [22].

The terrain is formed by numerous drainage channels, where the predominant soil type in both sites is the Dystrophic Red-Yellow Oxisol [23]. Characterized by deep soils, with low fertility and associated with ‘*terra firme*’ dense forests (not seasonally flooded). This soil type is a result of the decomposition of Precambrian rocks and, or in special cases, it results from tertiary sediments. Both areas are characterized by the presence of large trees such as ‘cupiúba’ (*Goupia glaba* Aubl. -Goupiaceae family), ‘matamatá’ [*Eschweilera odora* (Poepp. Ex O. Berg) -Lecythidaceae family], ‘envira preta’ (*Guatteria poepiggiana* Mart. -Malvaceae family) and ‘mandioqueira escamosa’ (*Qualea spp* -Vochysiaceae family). Some of these species can reach more than 50 m in height. The sites also have several species of palm trees and vines [24].

### Forest Inventory

The forest inventory was carried out by launching 100 random plots of 10 x 10 m in area A, and 10 systematically arranged plots of 10 x 10 m in area B (Fig S1). The systematic plots were arranged in a ‘fish bone’ pattern, keeping a distance of 50 m from each other. In area A, all trees with DBH (diameter at breast height, measured at 1.30 m from the ground) ≥ 40 cm were identified and their total biomass and height measurements measured. In total, the 180 trees recorded were distributed in the 82 species and 58 genera. Four plots (from 100) were drawn to quantify the forest biomass inventory of the smaller trees (5 ≤ DBH <40 cm). Area B had all trees with DBH ≥ 5 cm identified and their total height measurements were obtained with a TruePulse hypsometer (Vertex 360). DBH was always measured above the formation of the buttress root. All trees were felled to measure the total green biomass above the ground and the true total height (Ht).

### Allometric models

Six nonlinear models commonly used to describe the height-diameter relationship were tested (Table 1) [3,5,6,9,17]. Due to the low density of recorded trees per species and the absence of evident ecological gradients within the dataset, the models were fitted disregarding the species. The data set was randomly divided into two subsets: 70% (126 individual trees) for calibration and 30% (54 trees) for model validation. The DBH values for calibration ranged from 5.12 to 159.15 cm, and height ranged from 5.0 m to 51.0 m. The DBH in the validation set varied between 40.9 and 159 cm and the height between 22–51 m.

**Table 1.**
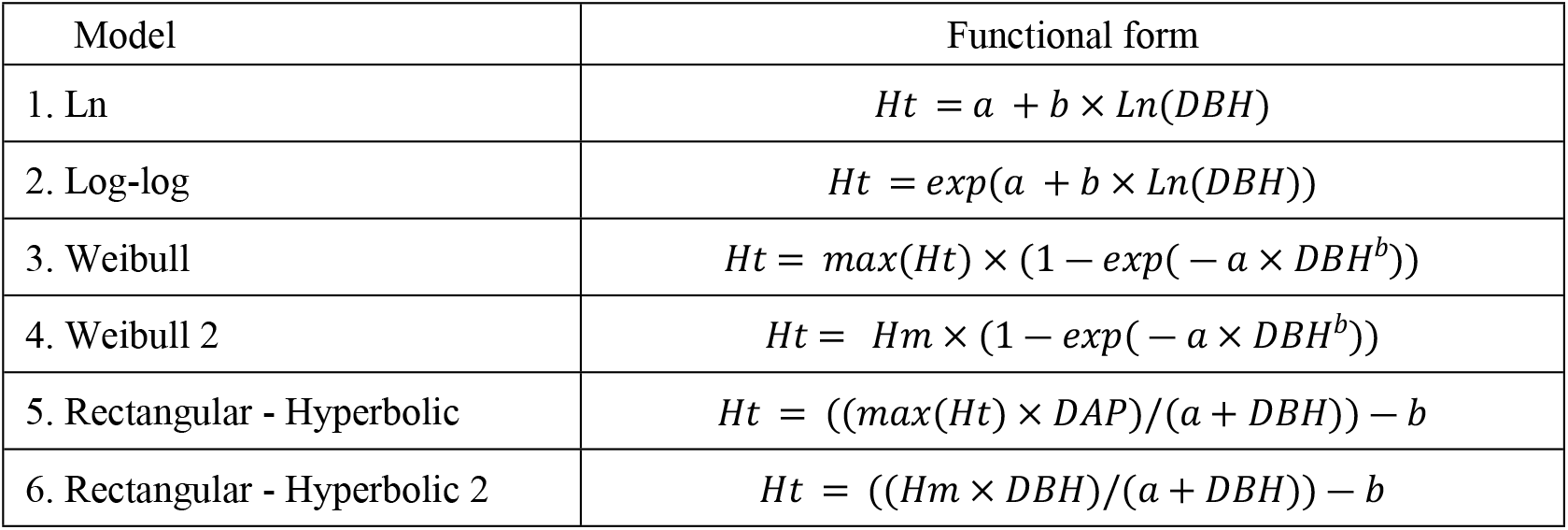
Hypsometric models used to describe the relationship between total height and diameter of the trees. a and b are the parameters to be estimated; DBH is diameter at breast height measured at 1.30 m from ground level in centimeters (cm); Hm is the maximum asymptotic total height of the sample in meters (m); max(Ht) is the highest total height of the sample, Ht is the total height of each tree in meters; ε is a random error; and Ln is the logarithm at the Neperian base (2.7128).

| Model | Functional form |
| --- | --- |
| 1. Ln | $H_t = a + b \times \ln(DBH)$ |
| 2. Log-log | $H_t = \exp(a + b \times \ln(DBH))$ |
| 3. Weibull | $H_t = \max(H_t) \times (1 - \exp(-a \times DBH^b))$ |
| 4. Weibull 2 | $H_t = H_m \times (1 - \exp(-a \times DBH^b))$ |
| 5. Rectangular - Hyperbolic | $H_t = ((\max(H_t) \times DBH)/(a + DBH)) - b$ |
| 6. Rectangular - Hyperbolic 2 | $H_t = ((H_m \times DBH)/(a + DBH)) - b$ |

We used the *nlme* package [25] in software R version 4.0.2 [26] to estimate the model parameters. The fitting was evaluated by the coefficient of determination (R²), root mean square error (RMSE), bias, maximum likelihood (AIC) and differences in AIC (Δ*i*) [27]. The Δ*i* was used to classify the models considering the empirical support proposed by [27]: Δ*i* ≤ 2 has substantial support, 2 < Δ*i* ≤ 10 has marginal support and Δ*i*> 10 has no support at all. For more details, see supplementary information (methods). The graphic analysis for the percentage residues was also carried out as a function of the estimated values of the height to identify models with homogeneous variance. Finally, the models were analyzed including the confidence limits (p <0.05) and regressing observed and estimated height compared to a linear trend of 1:1.

For the validation data (30% = 54 trees), in addition to the best local model, the individual height was estimated considering the regional equation proposed by [16], continental equation proposed by [6], and pantropical equation proposed by [17]. In addition, the estimated heights were also compared with the hypsometer measurements to analyze the accuracy and the error associated with the field measurements. The pantropical equation was based on the Weibull model, while the Continental equation for South America was based on the exponential form, and the regional equation was built based on log-log form for Santarém and Manaus data (see Table 2).

**Table 2.** Regional, continental and pan-tropical allometric equations tested to estimate total height. ht = total height; dbh = diameter at breast height (1,3 m ground level).

| Model | Funcional fit | Autor |
| --- | --- | --- |
| 1. Log-log | $Ht = 0.44 + 0.64 \times \ln(DBH)$ | [16] - Manaus |
| 2. Log-log | $Ht = 0.53 + 0.53 \times \ln(DBH)$ | [16] - Santarém |
| 3. Weibull | $Ht = \max(Ht) \times (1 - \exp(-0.5320 \times DBH^{0.5296})) + 0.007$ | [17] |
| 4. Exponential | $Ht = 35.83 - 31.15 \times \exp(-0.029 \times DBH)$ | [6] |

The performance of the models was evaluated comparing to real heights by the standard error (RSE) and the coefficient of variation (CV, p ≥ 0.05):

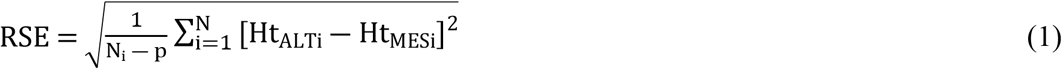

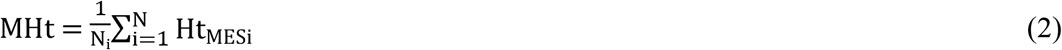

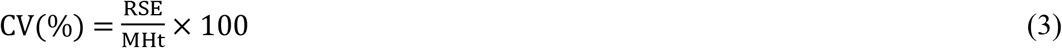

Where: Ht_ALTi_ and Ht_MESi_ are the estimated and observed height for tree i, respectively. The null hypothesis (H_0_) was that the heights estimated by one of the models or by hypsometer do not differ from the true height. When a significant effect was identified, the Tukey test (p < 0.05) was used to discriminate the differences between the groups. All analyzes were performed with Software R® version 4.0.2 (R Core Team 2020), using the *agricolae* package [28].

## Results

### Local allometric modeling

All the adjusted models had a determination coefficient greater than 84%, and the difference between the AIC values of the best and worst model was less than 10 units (Table 3). The model parameters were significant at a 95% significance level (p <0.05). The adjusted models showed good adjustments, producing satisfactory estimates concerning the trend of the observed data and valid confidence intervals (p <0.05) (Fig 2).

**Table 3.** Estimated parameters and model selection statistics for equations that describe the hypsometric relationship. The values in bold show the model better. R² is the correlation coefficient, AIC is the Akaike Information Criterion, Δi is the difference in AIC between each model and the best-fit model, wi is the Akaike weight, RMSE is the square root of the average error and the Bias is the systematic deviation of models.

| Model | Parameters |  |  | Performance fits |  |  |  |  |  | IC(< .05) |
| --- | --- | --- | --- | --- | --- | --- | --- | --- | --- | --- |
| | a | b | c | $R^2$ | AIC | $\Delta_i$ | $w_i$ | RMSE | Bias | |
| Ln | -10.0314 | 10.3848 |  | 0.840 | 926.15 | 7.24 | 0.027 | 4.369 | 0.049 | [16.67;24.96] |
| ln-ln | 1.5025 | 0.4808 |  | 0.844 | 921.59 | 2.68 | 0.262 | 4.307 | 0.076 | [16.77;24.99] |
| Rec | 46.5487 | -4.1623 |  | 0.845 | 920.37 | 1.46 | 0.482 | 4.291 | 0.062 | [16.71;24.92] |
| Rec2 | 53.3355 | -4.6631 | 53.4504 | 0.846 | 922.04 | 3.13 | 0.209 | 4.286 | 0.061 | [16.70;24.92] |
| **Weibull** | **0.062** | **0.6867** |  | **0.847** | **918.91** | **0** | **1** | **4.251** | **0.058** | [16.70;24.98] |
| Weibull2 | 0.0554 | 0.6215 | 64.5615 | 0.848 | 919.74 | 0.83 | 0.66 | 4.275 | 0.063 | [16.74;24.93] |

**Fig 2.**
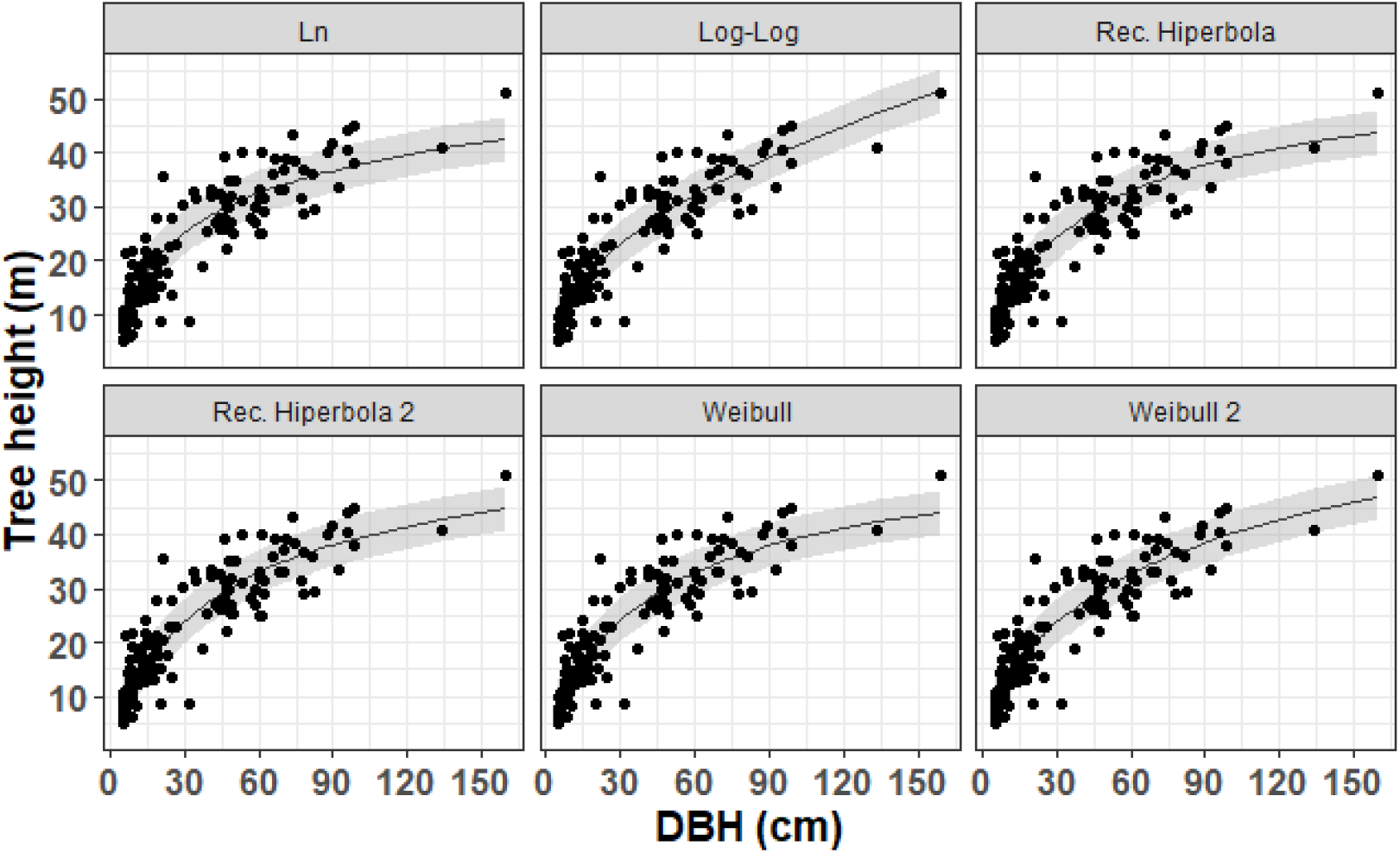
Estimates and prediction errors of local models adjusted for total height data in tropical forest in Amapá.

The best local model was the Weibull which showed higher accuracy, smaller error measures, smaller systematic deviation and higher support, with AIC = 918.910, R² = 0.84, RMSE = 4.251, Bias = 0.058, wi = 1.0 and Δi = 0 (Table 3 and Fig S2). The Weibull model 2 differed from Weibull mainly due to the asymptotic height. The visualization of the standardized residues according to the predicted height showed an absence of heteroscedasticity (Fig 3), concentrating the distribution and higher density of errors between -25 and 25%, which corroborates the fitting parameters (Table 3).

**Fig 3.**
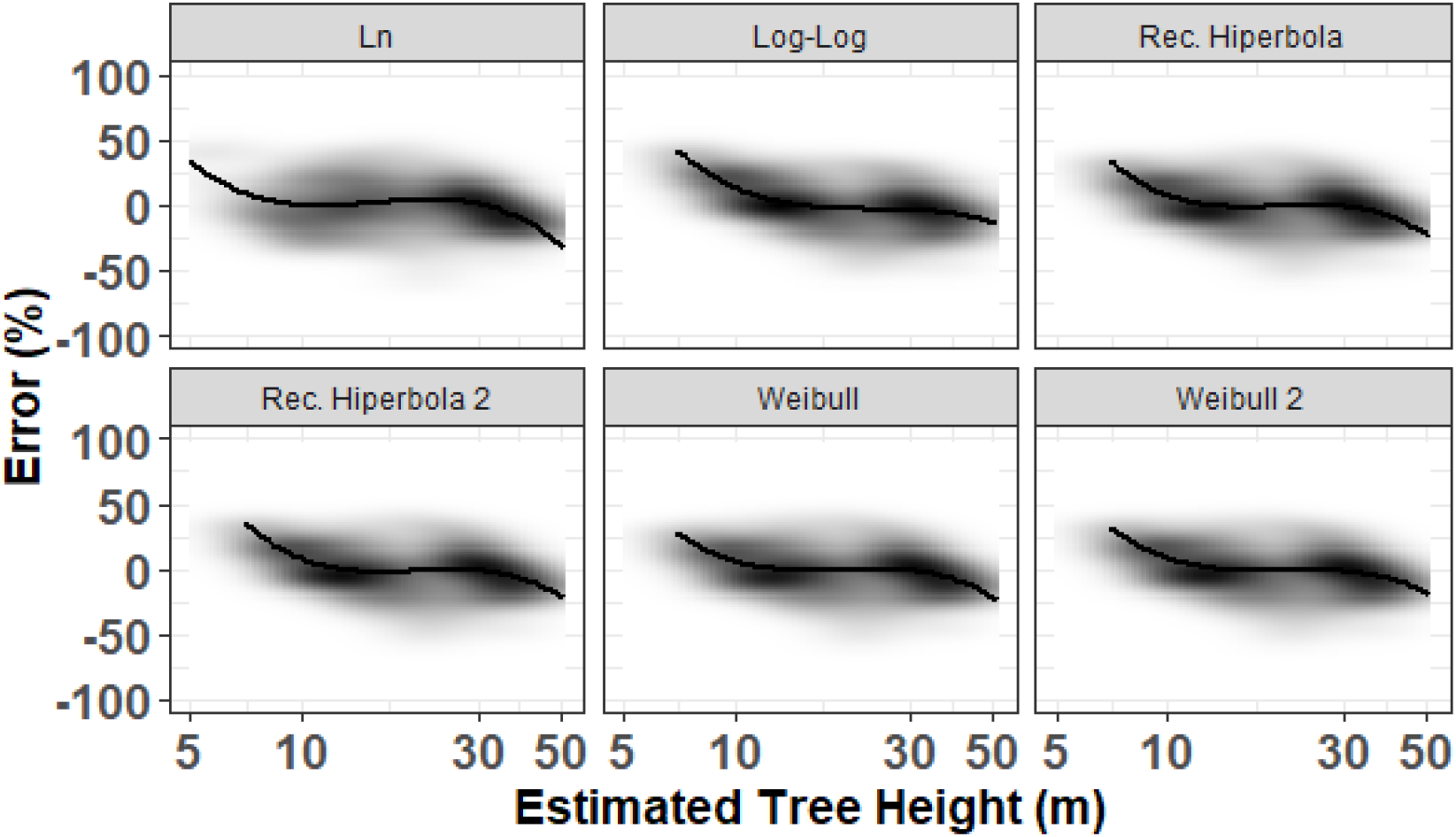
Percentage error (estimated Ht minus observed Ht, divided by the observed Ht, in%) vs estimated height values for the six non-linear theoretical models; the thick black line represents a spline regression of the data points, illustrating a slight negative bias at large estimated height values (values> 30 m). The background represents the density of the data point (n = 126 trees).

### Local, regional, continental and pantropical allometric models

The model comparison (best local adjusted, regional, continental and pantropical obtained from the literature) produced different trends, most of them differing at the total asymptotes. The validation dataset indicates that our local model adjusted in this study and the pantropical-Weibull model by [17] have similar precision and performance when predicting the tree heights. The continental model proposed by [6] and the regional models proposed by [16] underestimate height values when compared to the validation data (Fig 4). For those models, the relative error increased while height values also increased (Fig S3). The height estimated by the hypsometer showed large variation when compared to the validation dataset, but showed a consistent tendency to underestimate.

**Fig 4.**
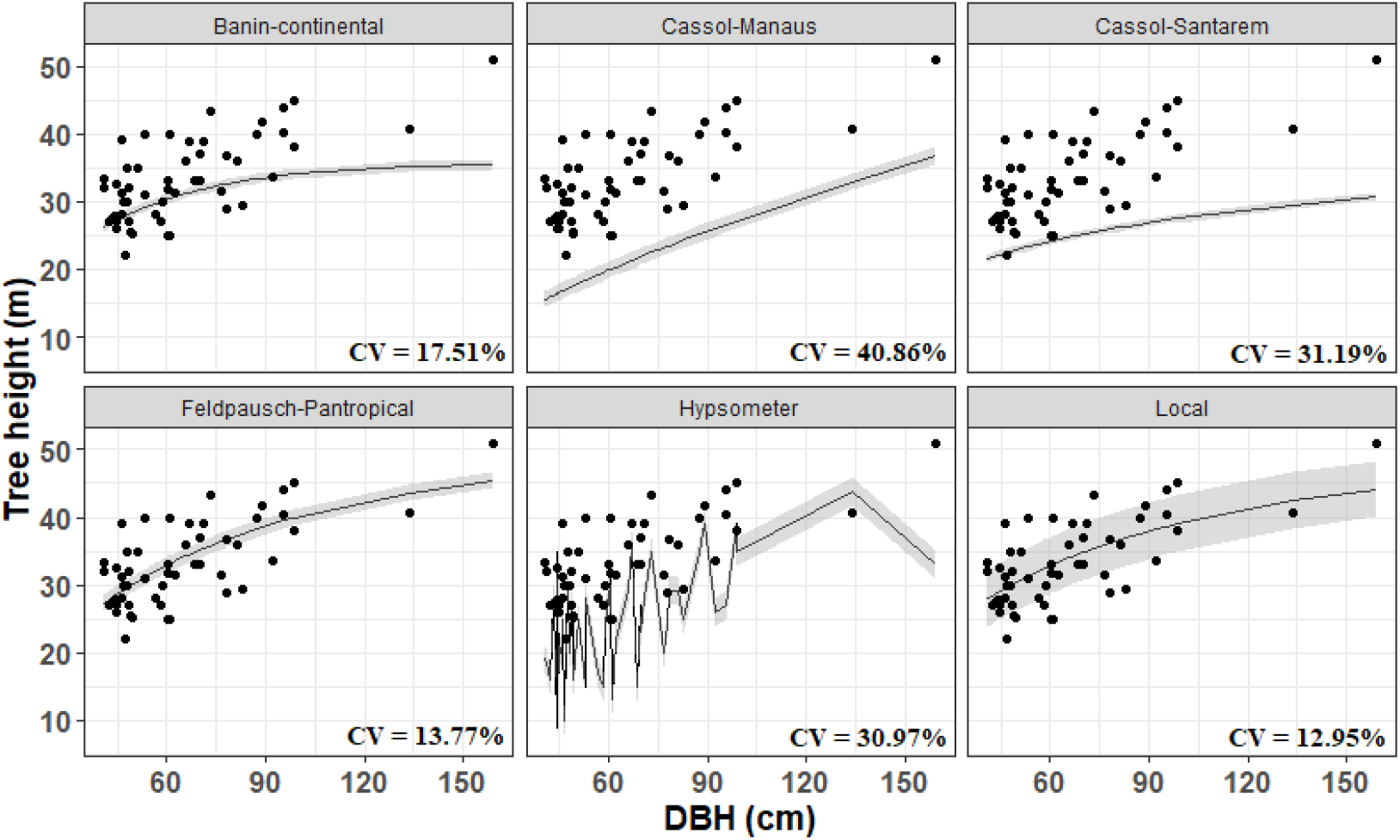
Estimates and percentage errors of allometric models (local, regional, continental and pantropical) and hypsometer for prediction of total height in tropical rainforest in Amapá.

The CV% practically doubles for regional-scale models and keeps stable for the continental and pantropical-scale models. The highest mean CV (40.86%) was noticed for the regional model proposed by [16] (Manaus regional model). The lowest mean bias was related to the local model (± 4.29 m), followed by the pan-tropical model by [17] (± 4.57m), both based on Weibull form. The local and pantropical models accurately estimated the average height (CV < 14%), showing low average forecasting errors (< 20%) even for the tallest trees (see Fig A3). Based on the statistical test, we can notice the superior performance of the local and pantropical models (Fstat = 60.92 p <. 001, df = 364, CV = 16.86%) (Fig 5, and also see table S2).

**Fig 5.**
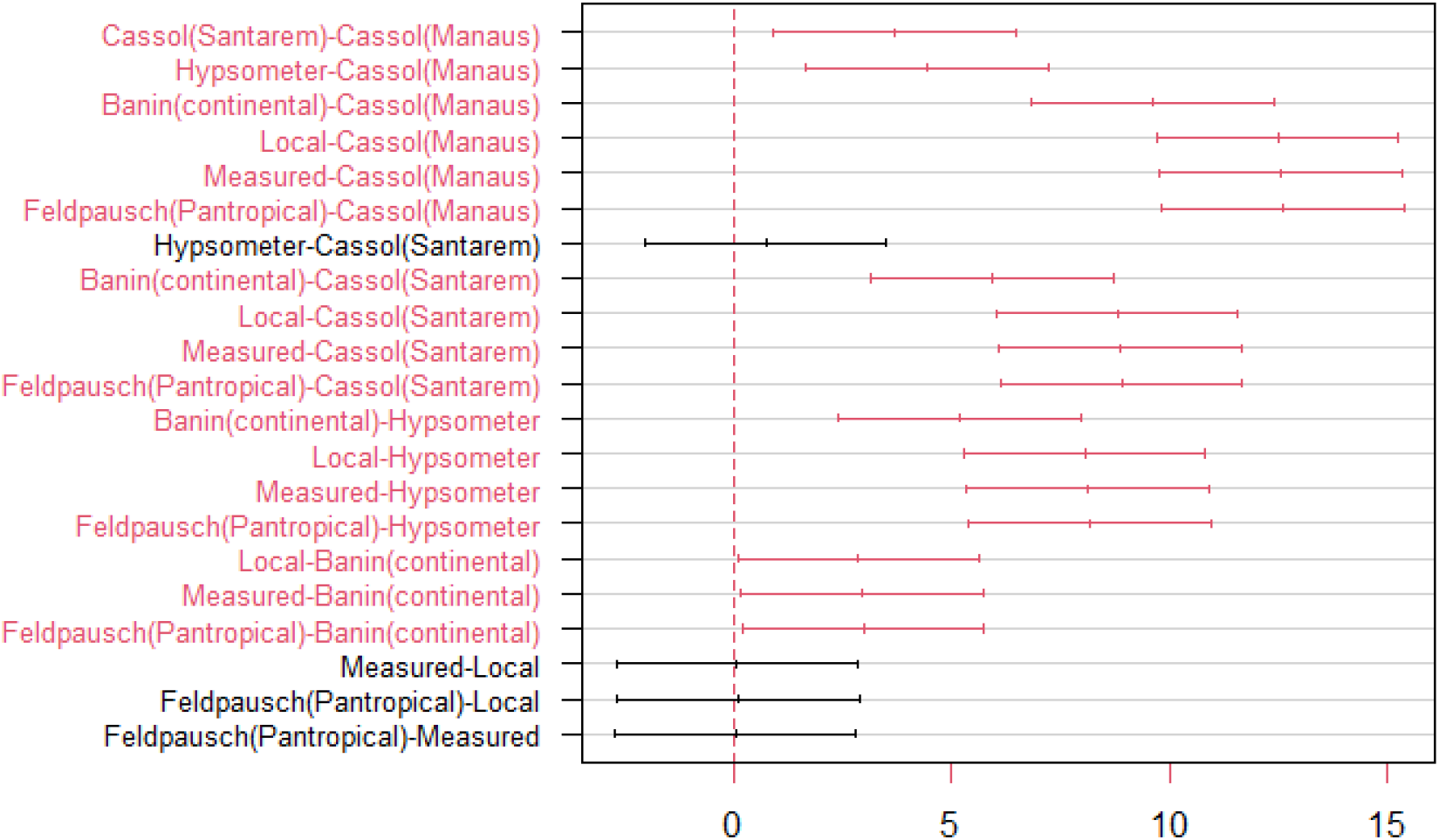
Intercomparison of average heights estimated by local, regional, continental and pantropical allometric models, by hypsometer and data measured in the field. The red contrasts indicate a statistical difference of zero by the Tukey test (p <0.05).

The tree height obtained by the hypsometer showed a systematic underestimation between diameter classes, when compared to the height recorded from the felled trees. We also noticed a larger amplitude for the tree height compared to observed range (Fig 6).

**Fig 6.**
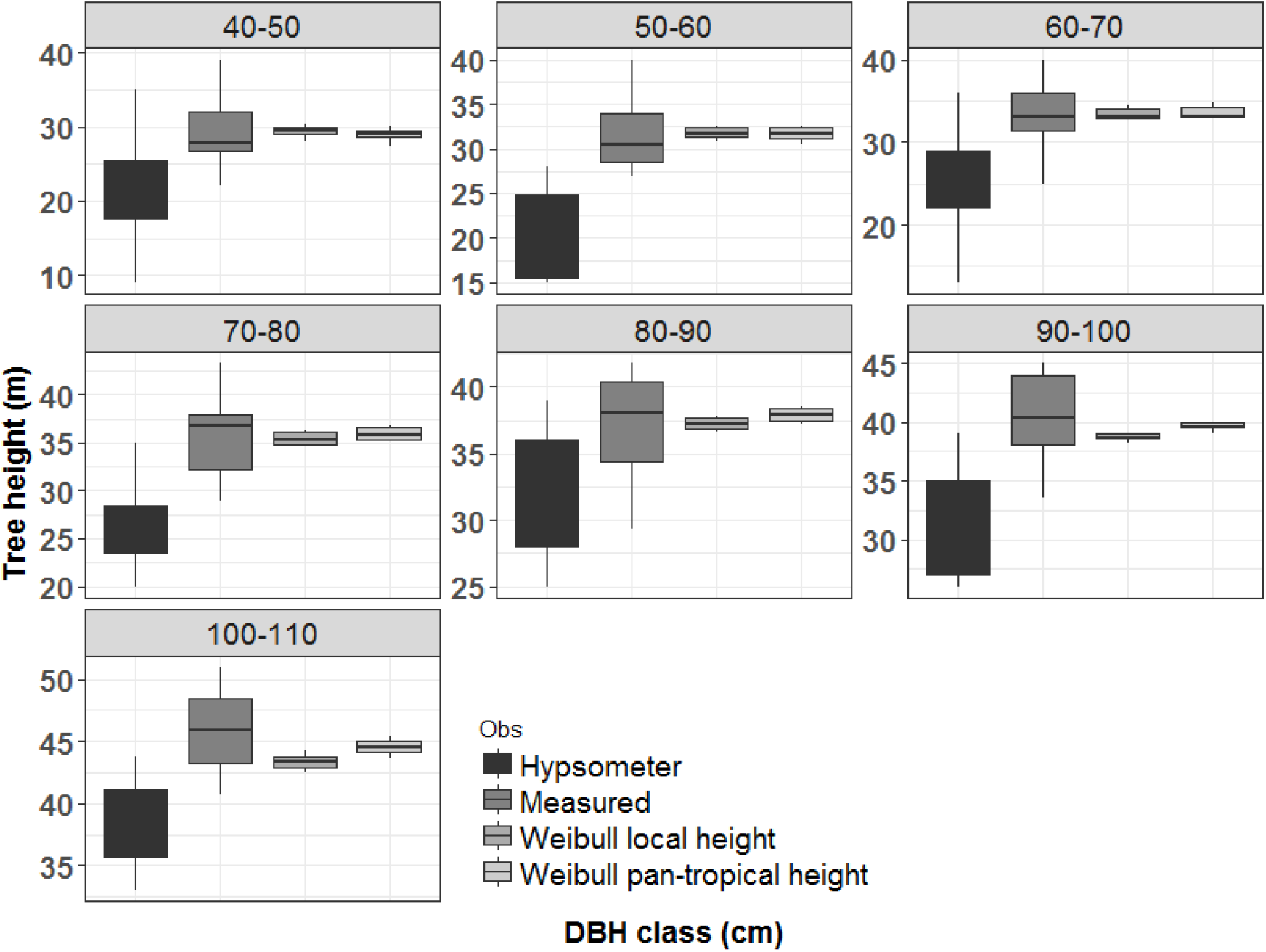
Comparison of height measurements by diameter class between the values generated by the hypsometer -Hypsometer, total length of the felled tree stem -Measured, height generated by the best local model -Weibull local height and height generated by the pan-tropical model – Weibull [17].

## Discussion

Tree height is essential to understand the dynamics of forest ecosystems, incorporating ecological characteristics of the species constituting the canopy. For example, the height can help to explain some of the carbon storage variations and productivity in ecosystem types [1]. Our results revealed a significant difference in the height recorded by the hypsometer when compared to the real height of felled trees. The errors produced by the hypsometer introduced large variation in heights in different diameter classes (see fig 6), indicating that the accuracy of this method depends on a strong trigonometric calibration and training [29]. On the other hand, it is necessary to assess the variability of these errors and possible trends in their occurrence, as well as to train field teams with the most up-to-date equipment and technological resources [2].

The tree height-diameter relationship depends on a series of physiological, biogeographic and environmental factors [7,30,31]. Besides climatic conditions and altitude [7,32,33] local edaphic factors, evolutionary and architectural restrictions [34,35], and the competition for space [36] could also influence the height-diameter relationship. [33] showed that in stressed environments by temperature and/or water the tree height is typically much lower than expected in a purely mechanical model.

In uneven forests, the great height variability within a diametric class may result in relative instability, resulting in measurement errors [37]. Local height-diameter allometric relationships implicitly incorporate geography and environment variation [7,17]. The allometric equation proposed in this work implements a local approach targeting the tropical rainforest from Northeast Amazonia (Amapá State). Which is characterized by the high heights variance and species heterogeneity, suggesting that a local approach should be recommended.

Most of the variation present in the height variable was explained by the local Weibull model (∼ 84%). The Weibull functional form is ecologically consistent and represents biological realism in dense forests [5,21]. The maximum height of the trees reaches a plateau through competition for light and nutrients [36] controlling the non-linear growth curve [5]. The correlation between diameter and height generates a form factor (trunk taper coefficient) that explains a good part of this behavior (r = 91%). The trunk shape variation also explains the growth in height, together with the size and shape of the crown [30,38,39].

Our results showed that the continental allometric models proposed by [6] and the regional model proposed by [16] are not suitable for dense forests in Amapá. Site properties may be affecting the tree height, resulting in interspecific and intraspecific variation [40,41]. We can find several examples in the literature about models that do not produce accurate estimates for some specific regions. For example, the pantropical height-diameter model by [17] did not produce accurate estimates of tree height in Asia [15]. In another study, regional models proposed for Central Africa [6,17] overestimated the tree heights for central Congo if compared to the pan-tropical model proposed by [7] [3,10].

Both regional models analyzed in our study showed a much higher average bias (+ 31.19%, and 40.86%, respectively Santarém and Manaus) when compared to the local model (+ 12.95%). A possible cause for the low performance is the low representativeness of the existent variations. Local allometric equations must consider the environmental gradients and species diversity to properly represent the variation presented in the vegetation [41]. This is exactly the limitation of the regional model, since they may not capture the existent heterogeneity observed for different scales [7,17]. [1] show that height-diameter models derived locally can be parameterized from 40 height observations. When the sampling is too small, the models may perform poorly in height prediction. The main reasons are: (i) non-linear relationships, such as asymptotic maximum heights, may not be evident in smaller sets of calibration data [42] and (ii) models can be excessively influenced by outliers (trees that are unusually tall or short for their diameter). [17] developed a set of regional-specific height-diameter allometries to minimize bias due to variation in canopy height observed in tropical regions (see also [6]).

In general, the deviations observed between models are associated with environmental variations [6] and species diversity [43–45]. The Amapá region is one of the most preserved locations, not only due to legal restrictions but also due to remoteness. Recent studies also evidenced that the resources availability and few disturbances allow plants to reach their potential in height [31]. The pantropical model presented accurate height estimates considering our validation dataset. We hypothesize that this result is explained by the similarity of the variation observed intercontinental and locally. Our results, in addition to confirming the accuracy of the best local model in different tree size classes, suggest that the generic pantropical model by [17] can be applied to the forests of Amapá.

## Conclusions

The local model proposed in this work performed similarly to the pantropical model by [17]. Establishing accurate height estimates requires a solid initial understanding of allometric models that explain or address the ecophysiological and environmental processes that govern vegetation dynamics and tree height growth. We suggest that future studies analyze the influence of different species and ecological gradients on the diameter/height ratio. The low performance of the hypsometer in height estimates reinforces the importance of incorporating new technologies in the process of measuring the attributes of individual trees such as LiDAR terrestrial or LiDAR aerial.

## Supporting information

Appendix A: Supplementary information for statistical analysis of nonlinear models adjusted to estimate total height in tropical rainforest in Amapá. Fig A1. Distribution of fixed sampling units of 10 x 10 m to carry out the forest inventory in Area B. Adapted from Oliveira et al. (2012). Appendix B: Supplementary information for the statistical results of adjustment and selection of non-linear models for total height estimation in tropical rainforest in Amapá. Fig A2. Predicted vs. scatter plot observed height (m) by the best local model (Weibull). The solid blue line represents the estimated values with gray confidence bands (p < 0.05). Red dashed lines represent predictions for height values. The solid black line represents the 1:1 ratio; Fig A3. Percent error (validation estimate minus observed Ht, divided by observed Ht, in %) vs estimated height values for the six height prediction alternatives; the thick black line represents a spline regression of the data points. The background represents the density of the data point (n = 54 trees); Table A1 – Comparison of mean height differences generated by local, regional, continental, pan-tropical and hypsometer model estimates with the true mean height measured in the field after tree thinning. Means followed by the same letters did not differ significantly by the Tukey test (p < 0.05).

## Author Contributions

Conceptualization, R.L. and E.G.; methodology, R.L., E.G., A.L.B.; R.C.L., E.S.; software, R.L. and E.G.; validation, R.L. and E.G.; formal analysis, R.L., E.G., F.E., J. A., C.O., A.L.B., D.S., A.P.B.; investigation,, R.L., E.G., F.E., J. A., C.O., A.L.B., D.S., A.P.B. and R.C.L., E.S.; resources,, R.L., E.G., R.C.L.; data curation, R.L. and R.C.L. writing—original draft preparation, R.L., E.G., F.E., J. A., C.O., A.L.B., D.S., A.P.B.; writing—review and editing, R.L., E.G. and F.E.; visualization, R.L. and E.G.; supervision, R.L. and E.G.; project administration, R.L. and R.C.L., E.S.; funding acquisition, R.L., E.G., R.C.L., E.S. All authors have read and agreed to the published version of the manuscript.

## Funding

Funding was provided by the Coordenação de Aperfeiçoamento de Pessoal de Nível Superior Brasil (CAPES; Finance Code 001); Conselho Nacional de Desenvolvimento Científico e Tecnológico (Processes 403297/2016-8 and 301661/2019-7) and (Processes 550467/2010-6)

## Data Availability Statement

Data can be made available upon request to authors.

## Acknowledgments

We thanks Universidade Federal dos Vales do Jequitinhonha e Mucuri (UFVJM) for support this work. We also thank the Universidade do Estado do Amapá (UEAP) for financial support to defray any fees for future publications of this manuscript.

## Conflicts of Interest

The authors declare no conflict of interest.

## Notes

### Competing Interest Statement

The authors have declared no competing interest.

